# iDeepLC: chemical structure information yields improved retention time prediction of peptides with unseen modifications

**DOI:** 10.1101/2025.10.31.685771

**Authors:** Alireza Nameni, Arthur Declercq, Ralf Gabriels, Robbe Devreese, Sven Degroeve, Lennart Martens, Robbin Bouwmeester

## Abstract

Deep learning has notably advanced the field of liquid chromatography–mass spectrometry-based proteomics. Accurate prediction of peptide retention times significantly enhances our ability to match LC-MS data with the correct peptides and proteins, especially for DIA data. While numerous models predict peptide LC retention times with high accuracy, few can accurately predict the retention times of chemically modified peptides, particularly those with modifications not encountered during model training.

In our previously developed DeepLC model, accurate predictions could be made for unseen modifications by leveraging the chemical composition of (modified) residues. Here, however, we present a further enhancement of this model based on chemical structural information. The resulting model, called iDeepLC, shows overall more accurate predictions, and better generalization performance for predicting the retention time of unseen modifications than DeepLC. iDeepLC is freely available as open-source software under the Apache2 license and can be found at https://github.com/CompOmics/iDeepLC.

## 1 INTRODUCTION

In Liquid Chromatography-Mass Spectrometry (LC-MS) proteomics, retention time refers to the specific duration it takes for a peptide to traverse the chromatography column and reach the MS detector. This value serves as an additional dimension that separates peptides prior to analysis, and thus substantially reduces sample complexity in the acquired MS dimensions. Deep learning models have been built that can accurately predict this expected retention time for a given peptide sequence under various experimental conditions. These predictions enable the *in-silico* creation of spectral libraries for the analysis of data-independent acquisition^1–4^ data, and enable the rescoring of peptide-spectrum-matches (PSMs) to increase identification sensitivity^5–9^. Furthermore, they are utilized to optimize experimental design^10^, to identify multiple peptides in chimeric fragmentation spectra^11^, to simulate LC-MS experiments^12^, and to provide for orthogonal validation of PSMs^13^.

However, accurate predictions of the retention time of modified peptides remains challenging. Most current predictors use a binary encoding for a limited set of possible modifications, where the model only recognizes the type and location of a specific modification in the sequence. Examples of such models include architectures such as transformers (Pham et al.)^14^, linear residual convolutional neural networks (CNNs) (Chronologer)^15^, capsule CNNs (DeepRT)^16^, neural networks with long short-term memory (LSTM) layers (Guan et al. and AutoRT)^17,18^, combinations of an LSTM and a transformer (DeepPhospho)^19^, and encoder-decoder models with gated recurrent units (Prosit)^20^. Only a few models employ a more sophisticated encoding method that includes the atomic composition of modifications, allowing the model to generalize over the modifications in the training set. This approach facilitates the prediction of peptides carrying modifications not observed during training, as seen in models using neural networks with LSTM layers (AlphaPeptDeep and pDeep3)^21,22^, and our own branched CNN, DeepLC^23^.

Here, we have explored the extension of these atomic count models by also providing structural information about the (modified) amino acid to the model. We hypothesized that this approach could further enhance the accuracy of retention time predictions for modified peptides and enable the differentiation of peptide isomers. Importantly, for small molecules, the issue of isomers has been addressed using Quantitative Structure-Selectivity Relationships (QSSR) and molecular descriptors^24^. Different sets of these molecular descriptor features have been used and compared in various machine learning models, with what is known as the MolLogP feature consistently emerging as the most discriminating^25^. This chemical descriptor represents the quantified propensity of a molecule to dissolve in non-polar solvents such as octanol, compared to polar solvents such as water ^26^.

Here we introduce iDeepLC short for ‘improved DeepLC’, a retention time predictor based on the DeepLC architecture, but that also incorporates structural and molecular information through the chemical descriptor MolLogP, in a branch neural network architecture with five paths. Our findings demonstrate that iDeepLC achieves superior predictive accuracy and generalization capabilities, making it a valuable tool for proteomics research.

## 2 METHODS

### 2.1 Model Architecture

iDeepLC employs a multi-branch neural network architecture similar to DeepLC^23^, with an additional, fifth branch that encodes chemical structural information. As explained in more detail in section 2.3, the first three branches consist of convolutional layers^27^ that encode chemical properties, diamino chemical properties, and atomic counts. A fourth branch is a fully connected layer that encodes general peptide features. The fifth branch consists of convolutional layers for a one-hot encoding of amino acids. The outputs of these five branches are then flattened and concatenated into several connected layers, finally yielding a single numerical output.

### 2.2 Incorporation of Additional Chemical Descriptor

To represent the structure of each, potentially modified, amino acid, we obtained their corresponding SMILES representation. This SMILES representation is then used to compute the MolLogP chemical structure descriptor with the RDKit library^28^ version 2023.3.2.

By incorporating MolLogP, iDeepLC gains insights into how atomic bonding patterns and individual atomic properties contribute to the hydrophobicity of amino acids. As a result, it can more effectively distinguish between various amino acids and their modifications, including those with identical atom compositions.

### 2.3 Input Encoding

The input of iDeepLC consists of, potentially modified, peptide sequences, each represented as a matrix with dimensions of 41 rows for the features described below, and 60 columns with each column representing a (modified) amino acid in a peptide with a maximum length of 60 amino acids. These features are categorized into five groups:

1. The **chemical descriptor, MolLogP**, is represented in the first row, encapsulating chemical characteristics of the peptides.
2. Rows two to seven encode the **atomic** counts of six elements: Carbon (C), Hydrogen (H), Nitrogen (N), Oxygen (O), Phosphorus (P), and Sulfur (S) present in the (modified) amino acids.
3. **Diamino chemical descriptor and diamino atoms** in rows eight and nine to fourteen, respectively. This branch sums the features of every two adjacent amino acids, enhancing the model’s ability to generalize across different peptide sequences. For peptides with an odd number of amino acids, the last column repeats the previous column’s features to ensure consistency without overlapping.
4. **Global features**, which spans rows 15 to 21, provides supplementary information about the peptide sequence. It includes the normalized length of the peptide relative to the maximum defined length, the atom composition of the first four and the last four amino acids, and the total atom composition of the entire peptide.
5. Rows 22 to 41 is the **One-hot encoding** to provide a clear, binary representation of amino acids within the sequence. This path is particularly important to distinguish between isoleucine and leucine as they are structural isomers^29^.

To address the challenge of varying peptides lengths, peptides with less than 60 amino acids are padded to maintain uniformity of 60 columns. This facilitates the straightforward encoding of various (modified) peptides regardless of their length. Note that in instances where modifications are present, the chemical descriptor, diamino chemical descriptor, and the atomic composition are adjusted to reflect modified amino acid change.

### 2.4 Hyperparameter tuning

The hyperparameters of the models were optimized on the Hela HF^30^ dataset. This optimization process was conducted with the WandB machine learning platform^31^. In total 18 different hyperparameters were tuned, these hyperparameter values are available in Supplementary Table 1. The tuned hyperparameters include the learning rate, batch size, dropout rate, kernel sizes, the number of channels, and the number of layers for the four convolutional layers.

### 2.5 Datasets and evaluations

We first evaluate performance of iDeepLC on 20 proteomics datasets (Supplementary Table 2), ranging from 3,000 to 160,000 sequences. These datasets mainly contain unmodified peptides and common modifications such as oxidation on methionine and carbamidomethyl on cysteine. One exception is ProteomeTools PTM^32^ dataset which includes synthetic peptides with various modifications. To ensure a fair and accurate comparison, we used the same training, testing, and validation sets that are used in the evaluation of DeepLC^23^.

Next, we assessed iDeepLC on 14 modifications from the ProteomeTools PTM dataset, an experiment referred to as the *14PTMs experiment* that was originally described in DeepLC. In this experiment we trained and optimized 14 iDeepLC models where each model only trained on peptides that did not contain a specific modification. These models were then evaluated on the remaining peptides, which all contained the modification excluded during training. We created two test sets from these remaining peptides to evaluate predictions: one where the excluded modification was encoded and one where it was ignored (not encoded) (Figure 1).

**Figure 1.**
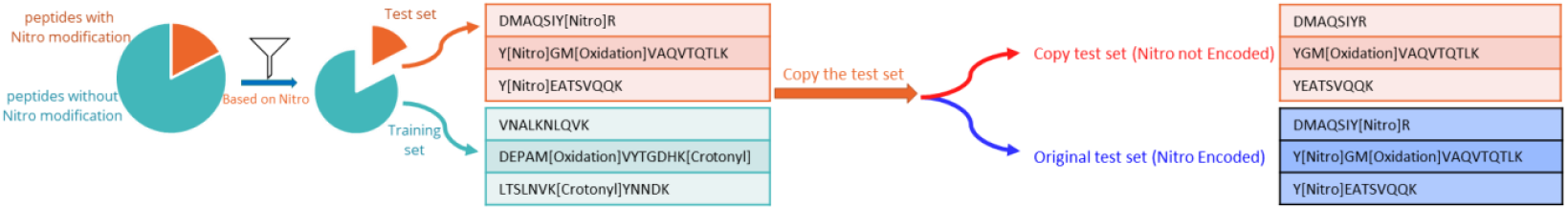
The process of generating datasets to evaluate the generalization of iDeepLC for peptides containing a specific peptide modification. In this example we filtered all peptides carrying a Nitro oxidation modification and trained the model on peptides without nitro oxidation and used those with the modification as a test set. In the next step the test set is duplicated, and all nitro oxidation modifications were removed (Nitro not Encoded) while in the other one we kept the peptides as they were (Nitro Encoded).

In an additional evaluation that we call the *glycine experiment*, we utilized all 20 datasets from Supplementary Table 2. We trained 19 models for each dataset, where each model excluded all peptides that contain a specific amino acid during training and was tested on peptides containing that amino acid. The excluded amino acids from training were initially represented as themselves and subsequently as glycine in a duplicated test set. Any modifications on the amino acids were disregarded when encoding it as glycine. This evaluation method was originally designed and described in DeepLC and was aimed at evaluating the generalization capabilities of iDeepLC. This evaluation assesses the model’s ability to predict the retention times of peptides incorporating amino acids absent from the training set. Furthermore, for 17 out of the 20 datasets there are many more training examples compared to the *14PTMs experiment* evaluation and should thus reflect actual performance better. Finally, for the comparison with DeepLC, we focused on two specific datasets: DIA HF^4^ which consists of approximately 113,000 sequences, and HeLa DeepRT^33^, with around 3,400 sequences.

For all evaluations three different metrics are used to assess performance: Mean Absolute Error (MAE), relative MAE, and the Pearson correlation. The relative MAE is used to make the comparison between different datasets possible. Equation (1) is used for the calculation of the relative MAE:

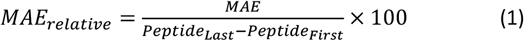

Here, the MAE is divided by the retention time difference between the first and last identified peptide in the respective dataset.

### 2.6 Training procedure

To initialize all models’ parameters for learning, we utilized the Kaiming uniform initialization^34^ method with a leaky ReLU^35^ activation function and we initialized biases to zero. This initialization scheme was applied to both convolutional and fully connected layers within the models. All layers use ELU^36^ as activation function except for fully connected layers that use ReLU^37^ activation function. The training parameters are available in Supplementary Table 1.

We trained the models for 1000 epochs for all 20 datasets and the *14PTMs evaluation*, respectively. For the *glycine evaluation*, we trained for either 300 or 1000 epochs, depending on whether the size of the training dataset was more than 1500. This decision to run fewer epochs is based on early convergence which was achieved much sooner for the larger training set sizes. In the training of all models, we saved the best model based on the MAE of the validation set. The training was done on an NVIDIA GeForce RTX 3090, using Python version 3.10.11 and PyTorch version 2.2.1 with CUDA version 11.8.

## 3 RESULTS

In this section, we first compare the performance of iDeepLC to DeepLC on 20 different datasets, followed by an evaluation of iDeepLC’s ability to generalize for (modified) peptides. This evaluation serves to assess its capability to generalize and is divided into two parts; first, iDeepLC’s capability to accurately predict the retention time of peptides with a specific modification excluded from training of the model (*14PTMs evaluation*), and second, the prediction for peptides containing an amino acid not used for training (*glycine evaluation*). It is expected that the addition of the MolLogP descriptor improves the ability of iDeepLC to generalize for modifications, especially for those unseen during training.

### 3.1 Prediction performance iDeepLC

We compared the prediction performance of iDeepLC with DeepLC on a set of 20 LC-MS datasets (see Methods). Figure 2 shows that iDeepLC demonstrates on-par performance with DeepLC. The largest difference in the relative MAE is observed for the ProteomeTools PTM dataset^32^, where the relative MAE differs more than 0.5%. The second biggest difference is for the Plasma 1h datasets with a much smaller 0.28% relative MAE difference. It is important to emphasize that both iDeepLC and DeepLC were trained, validated, and evaluated using identical data splits across each dataset.

**Figure 2.**
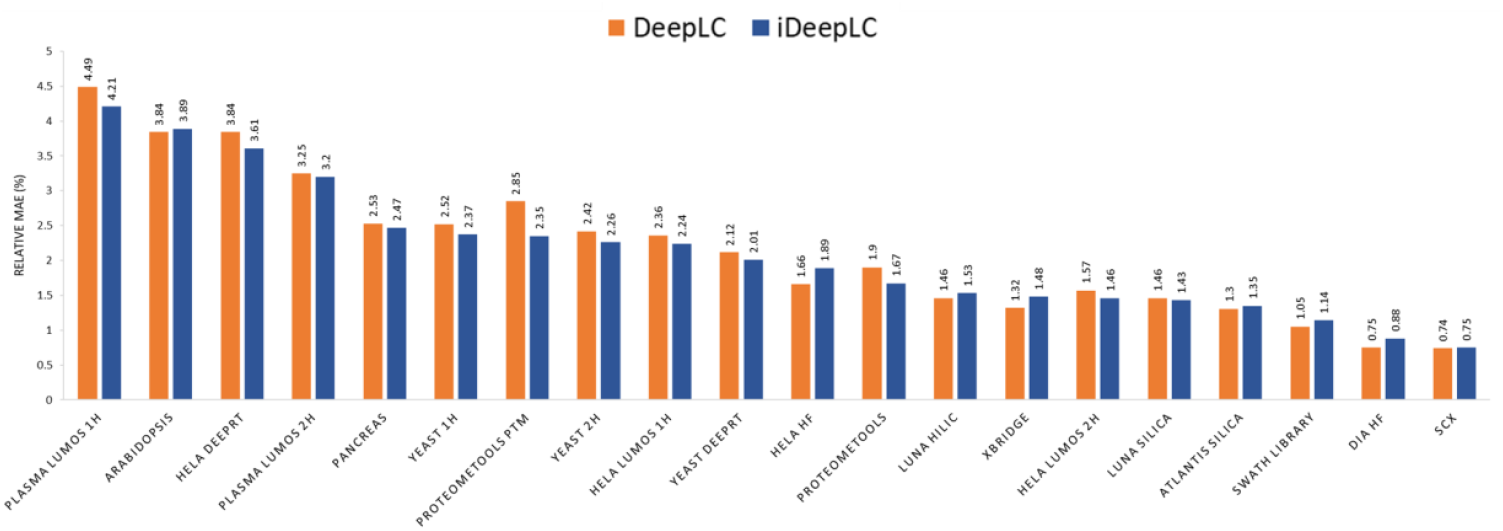
Performance comparison of DeepLC (orange) and iDeepLC (blue) for 20 datasets with the MAE as metric (lower is better).

### 3.2 PTM evaluation

The improved performance on the dataset consisting primarily of modified peptides (ProteomeTools PTM dataset) indicates that iDeepLC can generalize more effectively for modified peptides. Due to the additional chemical descriptor, iDeepLC should be able understand the peptides’ physicochemical properties better compared to DeepLC, where modified peptides are only represented with their atomic composition. This improvement of iDeepLC to predict for modified peptides is further investigated in the *14PTMs evaluation*. In this evaluation the Proteome Tools PTM ^32^ is used to assess how well iDeepLC understands the modified peptides’ physicochemical properties, without explicitly training on observations for a specific amino acid modification.

Figure 3 highlights two modifications, nitro oxidation and deamidation, from the *14PTMs evaluation*. For these modifications, iDeepLC can predict the modification induced retention time shift much better than DeepLC. For these two modifications, the MAE for deamidation and nitro oxidation are improved with 58% (from 257 to 109 seconds) and 52% (from 567 to 272 seconds), respectively. For completeness, the scatter plots of all modifications are available in Supplementary Figure 1.

**Figure 3.**
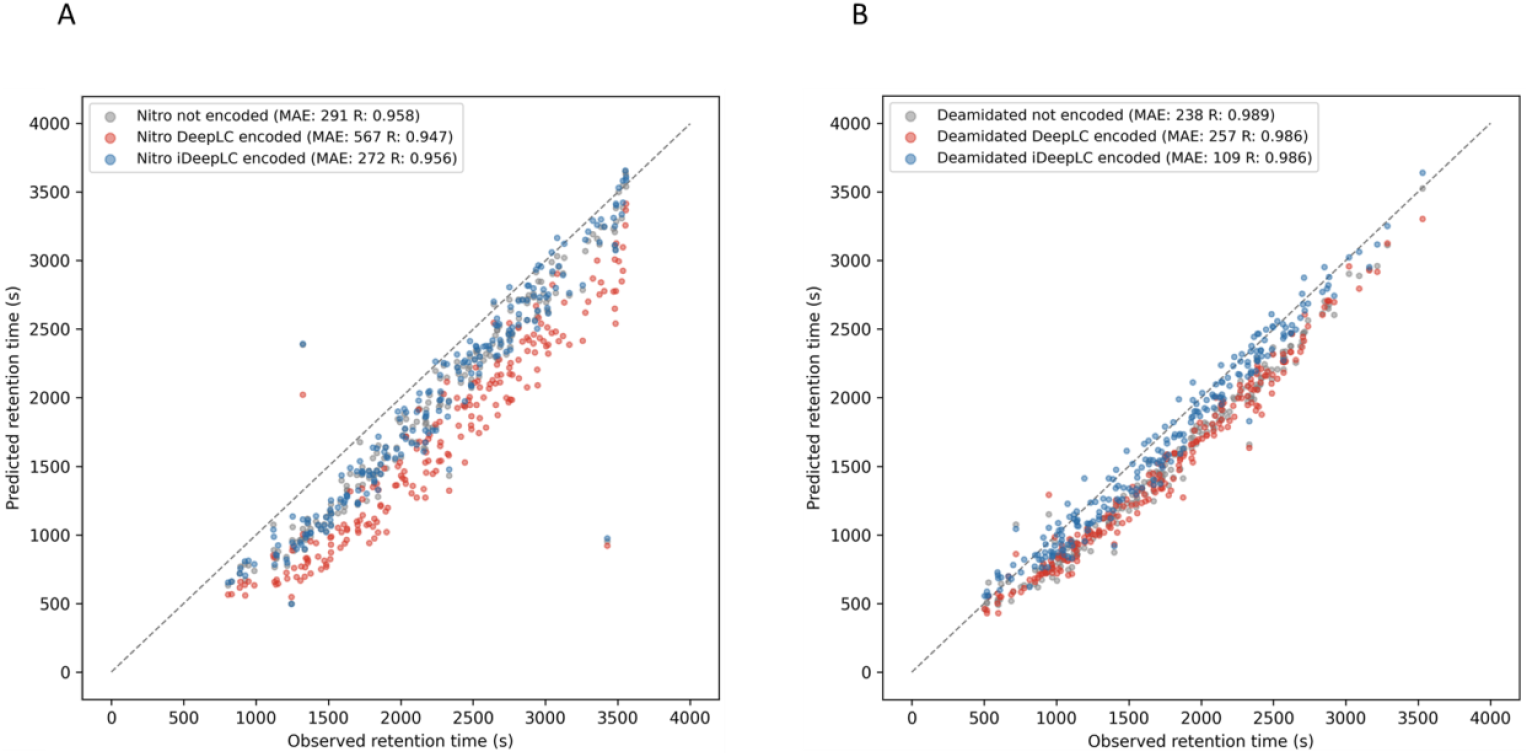
Performance comparison of DeepLC and iDeepLC for models that were not trained on peptides containing nitro oxidation,**A** and deamidation,B but evaluated on their respective modifications. The blue dots and the red dots show the retention time predicted by iDeepLC and DeepLC respectively tested with encoded modifications, and the grey dots show the retention time on an iDeepLC model tested without encoding modifications.

This trend of better performance for iDeepLC and DeepLC over the baseline continues for the remaining twelve modifications, where performance is either better or comparable (Figure 4). Furthermore, these results show that the impact of encoding or not encoding for the methyl group modifications is minimal. In contrast, there is a large difference between encoding and not encoding for acetyl, succinyl, propionyl, crotonyl, malonyl, oxidation and deamidation. Also, these modifications are much more accurately predicted when encoded for both prediction models. The sole exception are the phosphorylated peptides which are more accurately predicted when not encoded compared to when encoded. However, the prediction error of iDeepLC is still smaller than DeepLC when encoding the phosphorylation. This improved prediction accuracy of iDeepLC over DeepLC is also noticeably clear for carbamidomethylation, deamidation, oxidation, and nitro oxidation.

**Figure 4.**
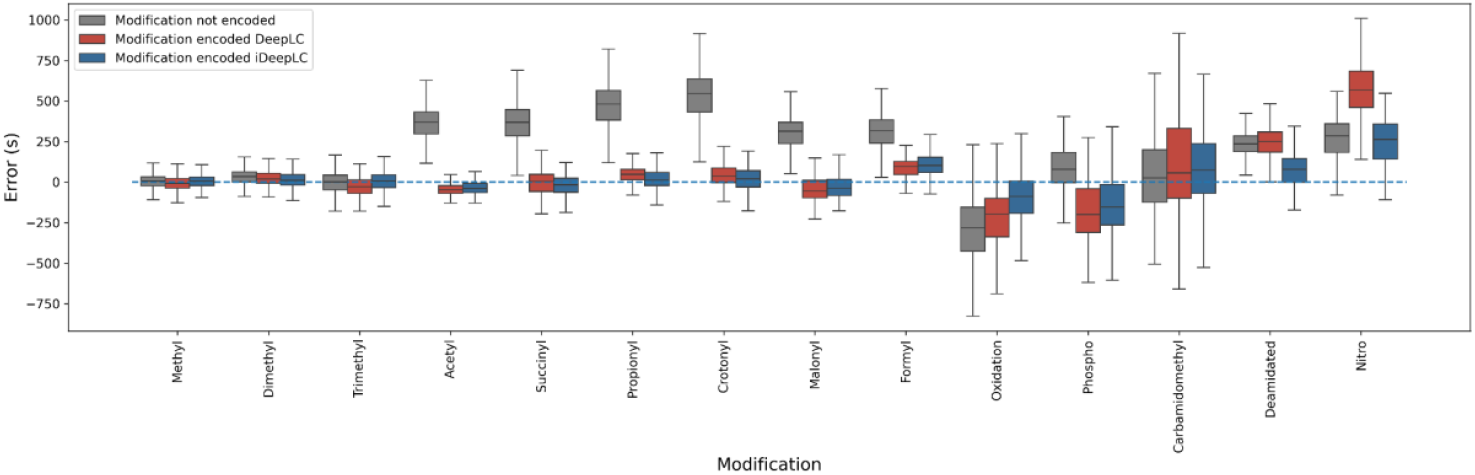
Each modification used for testing is shown on horizontal axis with its corresponding error of predicted and observed retention time. Each modification has three boxes, when the evaluated modification was not encoded (grey) or encoded and predicted by DeepLC (red) or encoded and predicted by iDeepLC (blue).

The relative differences with the not encoded baseline and DeepLC or iDeepLC are used to further investigate performance differences between these models (Figure 5 and Supplementary Table 3). In figure 5 any modifications positioned above the diagonal line indicate a superior performance for iDeepLC. Any point that is below the diagonal means better performance for DeepLC. As observed before, iDeepLC outperforms DeepLC in predicting the retention times for phosphorylation, carbamidomethylation, deamidation, oxidation, and nitro oxidation modifications. For the remaining nine modifications the performance of iDeepLC is on par with DeepLC.

**Figure 5.**
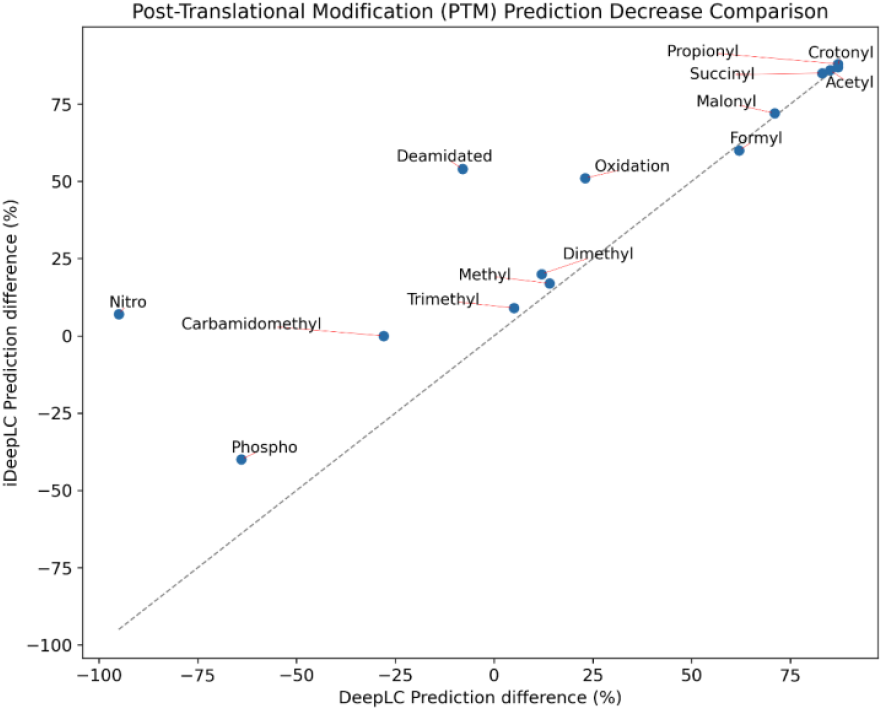
iDeepLC and DeepLC percentage difference in MAE compared to the not encoded test set (baseline) for each modifications in the 14PTMs evaluation.

### 3.3 Modified glycine evaluation

In the *glycine evaluation*, the same methodology as the *14PTMs evaluation* is followed, but instead of modifications, amino acids are excluded from training and tested on. Furthermore, instead of not encoding the amino acid, a baseline is created by replacing the specific amino acid with glycine.

As expected, this *glycine evaluation* shows that encoding amino acids as themselves results in a lower MAE in most cases (Figure 6). This improved performance can be seen for the individual amino acids in Figure 6 positioned above the diagonal line, where the distance to the diagonal line quantifies this improvement over the baseline where amino acids are encoded as glycine. This also means that amino acids below the diagonal line have the reverse conclusion, encoding the amino acid as glycine performs better. Notably, the largest improvements over the baseline are achieved for hydrophobic amino acids. Indeed, correctly encoding amino acids that have the largest contribution to a peptide’s hydrophobicity, and thus retention time in reversed phase chromatography, greatly impacts the prediction accuracy. Importantly, this large hydrophobic contribution of specific amino acids is very effectively captured in the iDeepLC model. iDeepLC does show slightly worse performance for amino acids like lysine (K), but this is likely due to its distinct polar properties and small training dataset size after excluding tryptic peptides.

**Figure 6.**
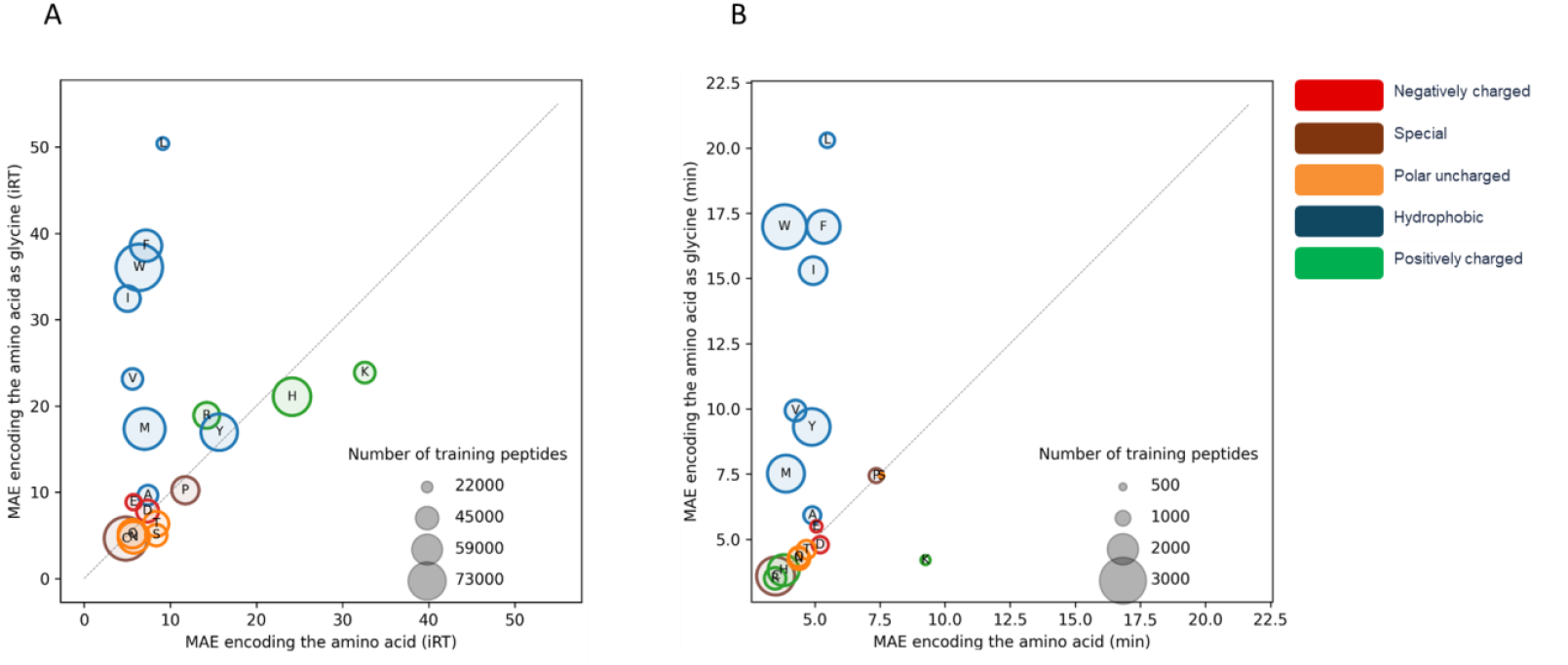
Glycine evaluation where each amino acid that was excluded is shown in a circle with its size determining the number of training peptides and its color shows the chemical property. An amino acid’s position indicates the MAE for all peptides containing that amino acid and it is either encoded as glycine (vertical axis) or as its own atomic composition (horizontal axis). Everything above the diagonal line is predicted with a higher accuracy when the amino acid is encoded as itself. Two datasets are used, **A**, DIA HF (bigger dataset), **B**, HeLa DeepRT (smaller dataset)

We summarized the results of the *glycine evaluation* for 15 datasets with a reverse-phase stationary phase in Figure 7. This figure shows the relative MAE for each amino acid in all reverse-phased datasets when encoded as themselves or when encoded as glycine. Among the 19 amino acids, encoding the amino acid as itself showed improvements for eleven amino acids (A, E, F, I, L, M, P, R, V, W, and Y). For three amino acids (C, H, and K), the results for encoding the amino acids were worse compared to the baseline. Finally, for the remaining five amino acids (D, N, Q, S, T) there is a minimal difference between encoding them as their respective amino acid and encoding them as glycine.

**Figure 7.**
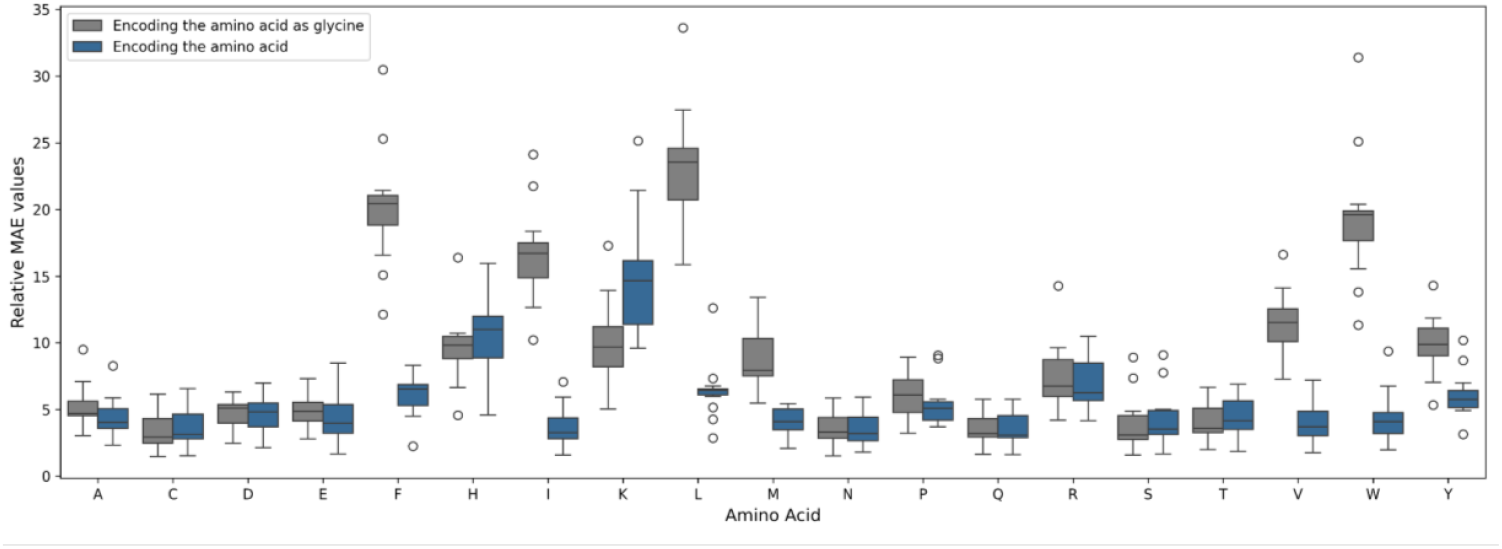
Relative MAE for each amino acid across all fifteen reverse-phased datasets. The boxplots illustrate the comparison between encoding the amino acid as itself (blue) versus encoding it as glycine (gray).

Instead of comparing iDeepLC to a baseline here, the model was also compared to DeepLC with the same *glycine evaluation* (Figure 8). In the DIA HF dataset, iDeepLC has higher prediction accuracy in 12 out of 19 cases, while for the much smaller dataset, HeLa DeepRT a lower MAE was achieved in 14 out of 19 cases. Notably, in both datasets most of the hydrophobic amino acids such as tryptophan (W), are much better predicted by iDeepLC. Furthermore, for the cases where iDeepLC is outperformed by DeepLC, there is only a small gain in prediction accuracy.

**Figure 8.**
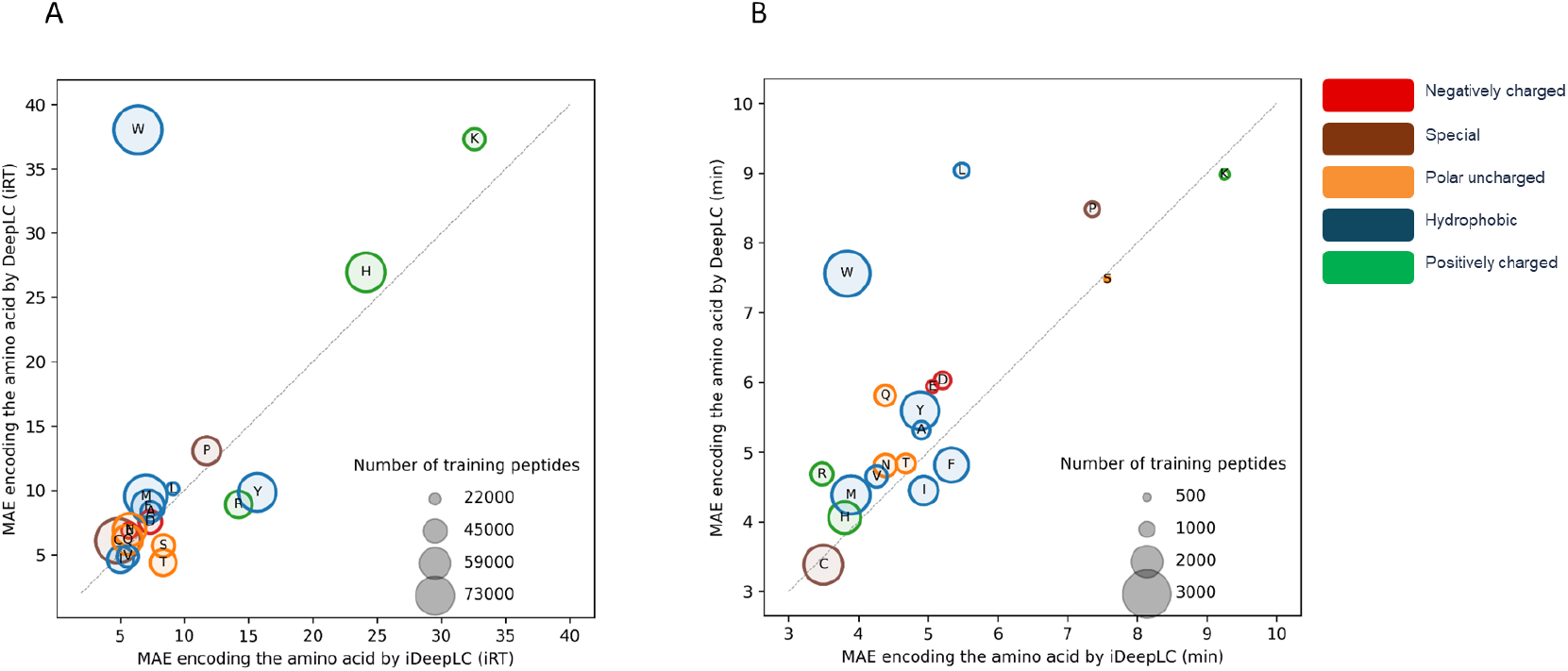
Glycine evaluation where each amino acid that was excluded is shown in a circle with its size determining the number of training peptides and its color shows the chemical property of that amino acid. These figures compare the error of predicted and actual retention time between DeepLC (vertical axis) and iDeepLC (horizontal axis). Everything above the diagonal line shows the better performance of iDeepLC compared with DeepLC. Two datasets are used, **A**, DIA HF (bigger dataset), **B**, HeLa DeepRT (smaller dataset)

## 4 CONCLUSION & DISCUSSION

Our results show that adding the MolLogP descriptor enables iDeepLC to learn the physicochemical characteristics of modifications. This understanding means that their impact on the LC retention time is more accurately predicted. Notably, iDeepLC delivers state-of-the-art predictive accuracy for unmodified peptides, while also exhibiting superior performance when applied to modified peptides. This was highlighted in two evaluations where iDeepLC outperforms DeepLC for specific modifications and predicting retention times for previously unobserved amino acids.

The only drawback for iDeepLC is that it requires the structure of amino acids and their modifications to obtain the chemical descriptor. However, this is readily resolved in the future as these LC-MS behavior predictors are gaining traction and the value of incorporating structural information as input to these models is more apparent.

## 5 DATA AVAILABILITY

All data used to train and evaluate iDeepLC are available on Zenodo at https://doi.org/10.5281/zenodo.15011301 and it contains the following projects: HeLa hf^30^, ProteomeTools^38^, SWATH library^39^, Plasma lumos 1h^40^, DIA HF^4^, HeLa lumos 2h^40^, Pancreas^41^, Xbridge^42^, ATLANTIS SILICA^42^, LUNA SILICA^42^, LUNA HILIC^42^, SCX^42^, Yeast 2h^43^, HeLa lumos 1h^40^, Yeast 1h^43^, Arabidopsis^44^, Yeast DeepRT^45^, ProteomeTools PTM^32^, Plasma lumos 2h^40^, and HeLa DeepRT^33^.

## 6 ACKNOWLEDGEMENTS

A.N. acknowledges funding from the European Union’s Horizon 2020 research and innovation programme under the Marie Skłodowska-Curie grant agreement N° 956148. R.D., A.D., R.G., L.M. and R.B. acknowledge funding from the Research Foundation Flanders (FWO) [1SH9O24N, 12B7123N, 1SE3724N, G010023N, G028821N, 12A6L24N]. L.M. acknowledge funding from the Horizon Europe Projects BAXERNA 2.0 [101080544] and COMBINE [101191739], and from the Ghent University Concerted Research Action [BOF21/GOA/033]. L.M. is further supported by the CHIST-ERA project ODEEP-EU [G0GDV23N] and F.I. by Ghent University Starting Grant BOF/STA/202209/011.

