## Supplementary document for "iDeepLC: chemical structure information yields improved retention time prediction of peptides with unseen modifications"

Supplementary Table 1: Distribution of hyperparameters values

| **Hyperparameters** | **Distribution** | **Final value** |
| --- | --- | --- |
| Batch size | [min:64, max:512] | 96 |
| Dropout | [min:0.05, max:0.6] | 0.23 |
| CNN1 channels | [min:32, max:512] | 245 |
| CNN1 layers | [min:1, max:10] | 1 |
| CNN1 kernel size | [3, 5, 7, 9, 11, 13, 15, 17] | 5 |
| CNN2 channels | [min:16, max:256] | 41 |
| CNN2 layers | [min:0, max:7] | 0 |
| CNN2 kernel size | 3, 5, 7, 9, 11, 13] | 3 |
| CNN3 channels | [min:32, max:512] | 35 |
| CNN3 layers | [min:1, max:10] | 5 |
| CNN3 kernel size | [3, 5, 7, 9, 11, 13, 15, 17] | 9 |
| CNN4 channels | [min:4, max:64] | 50 |
| CNN4 layers | [min:0, max:4] | 3 |
| CNN4 kernel size | [3, 5, 7, 9, 11, 13] | 7 |
| Fully connected1 output | [min:8, max:256] | 78 |
| Fully connected1 layers | [min:1, max:8] | 2 |
| Fully connected2 output | [min:8, max:256] | 77 |
| Fully connected2 layers | [min:1, max:8] | 1 |

Supplementary Table 2: Datasets used in iDeepLC

| Name | Data origin for fitting | Training peptides | Validation peptides | Test peptides | Repository identifier | Downloaded raw file name pattern | Samples (including replicates) | Fractions per sample | Runs | Column type |
| --- | --- | --- | --- | --- | --- | --- | --- | --- | --- | --- |
| HeLa HF | Custom workflow | 137821 | 7253 | 16119 | PXD006932 | 46frac_15min_15000 | 1 | 46 | 46 | RP |
| ProteomeTools | Custom workflow | 125331 | 6596 | 14658 | PXD010595; PXD004732 | - | - | - | - | RP |
| SWATH library | DeepRT4 | 125331 | 6596 | 14658 | PXD000954 | - | - | - | - | RP |
| Plasma Lumos 1h | Custom workflow | 2997 | 157 | 350 | PXD013477 | plasma_DDA_1h | 2 | 1 | 2 | RP |
| DIA HF | Guan et al.7 | 96798 | 5094 | 11321 | PXD005573 | - | - | - | - | RP |
| HeLa Lumos 2h | Custom workflow | 49495 | 2604 | 5788 | PXD013477 | Hela_DDA_2h | 2 | 1 | 2 | RP |
| Pancreas | Custom workflow | 43002 | 2263 | 5029 | PXD010154 | P013678 | 36 | 1 | 36 | RP |
| Xbridge | DeepRT4 | 34231 | 1801 | 4003 | - | - | - | - | - | HILIC |
| ATLANTIS SILICA | DeepRT4 | 33421 | 1759 | 3909 | - | - | - | - | - | HILIC |
| LUNA SILICA | DeepRT4 | 31483 | 1656 | 3682 | - | - | - | - | - | HILIC |
| LUNA HILIC | DeepRT4 | 30848 | 1623 | 3607 | - | - | - | - | - | HILIC |
| SCX | DeepRT4 | 26051 | 1371 | 3047 | - | - | - | - | - | SCX |
| Yeast 2h | Custom workflow | 23512 | 1237 | 2750 | PXD003472 | 500ng_120min | 4 | 1 | 4 | RP |
| HeLa Lumos 1h | Custom workflow | 21638 | 1138 | 2530 | PXD013477 | Hela_DDA_1h | 2 | 1 | 2 | RP |
| Yeast 1h | Custom workflow | 15822 | 832 | 1850 | PXD003472 | 500ng_60min | 4 | 1 | 4 | RP |
| Arabidopsis | Custom workflow | 13310 | 700 | 1556 | PXD008812 | R1 | 20 | 1 | 20 | RP |
| Yeast DeepRT | DeepRT4 | 12197 | 641 | 1426 | - | - | - | - | - | RP |
| ProteomeTools PTM | Custom workflow | 4867 | 256 | 569 | PXD009449 | - | - | - | - | RP |
| Plasma Lumos 2h | Custom workflow | 3659 | 192 | 428 | PXD013477 | plasma_DDA_2h | 2 | 1 | 2 | RP |
| HeLa DeepRT | DeepRT4 | 2917 | 153 | 341 | - | - | - | - | - | RP |

Supplementary Table 3: MAE comparison of iDeepLC and DeepLC on encoded test sets, highlighting the percentage difference from the not encoded test set (baseline)

| **PTM** | **baseline** | **DeepLC** | **iDeepLC** | **DeepLC decrease (%)** | **iDeepLC decrease (%)** | **difference increase** |
| --- | --- | --- | --- | --- | --- | --- |
| Methyl | 59 | 51 | 49 | 13.56 | 16.95 | 3.39 |
| Dimethyl | 59 | 52 | 47 | 11.86 | 20.34 | 8.47 |
| Trimethyl | 64 | 61 | 59 | 4.69 | 7.81 | 3.12 |
| Acetyl | 364 | 54 | 51 | 85.16 | 85.99 | 0.82 |
| Succinyl | 363 | 62 | 55 | 82.92 | 84.85 | 1.93 |
| Propionyl | 476 | 64 | 57 | 86.55 | 88.03 | 1.47 |
| Crotonyl | 539 | 70 | 69 | 87.02 | 87.20 | 0.18 |
| Malonyl | 327 | 96 | 92 | 70.64 | 71.88 | 1.24 |
| Formyl | 312 | 117 | 124 | 62.50 | 60.26 | -2.24 |
| Oxidation | 316 | 244 | 155 | 22.78 | 50.95 | 28.16 |
| Phospho | 138 | 226 | 193 | -63.77 | -39.86 | 23.91 |
| Carbamidomethyl | 208 | 267 | 209 | -28.37 | -0.48 | 27.89 |
| Deamidated | 238 | 257 | 109 | -7.98 | 54.20 | 62.18 |
| Nitro | 291 | 567 | 272 | -94.85 | 6.53 | 101.38 |

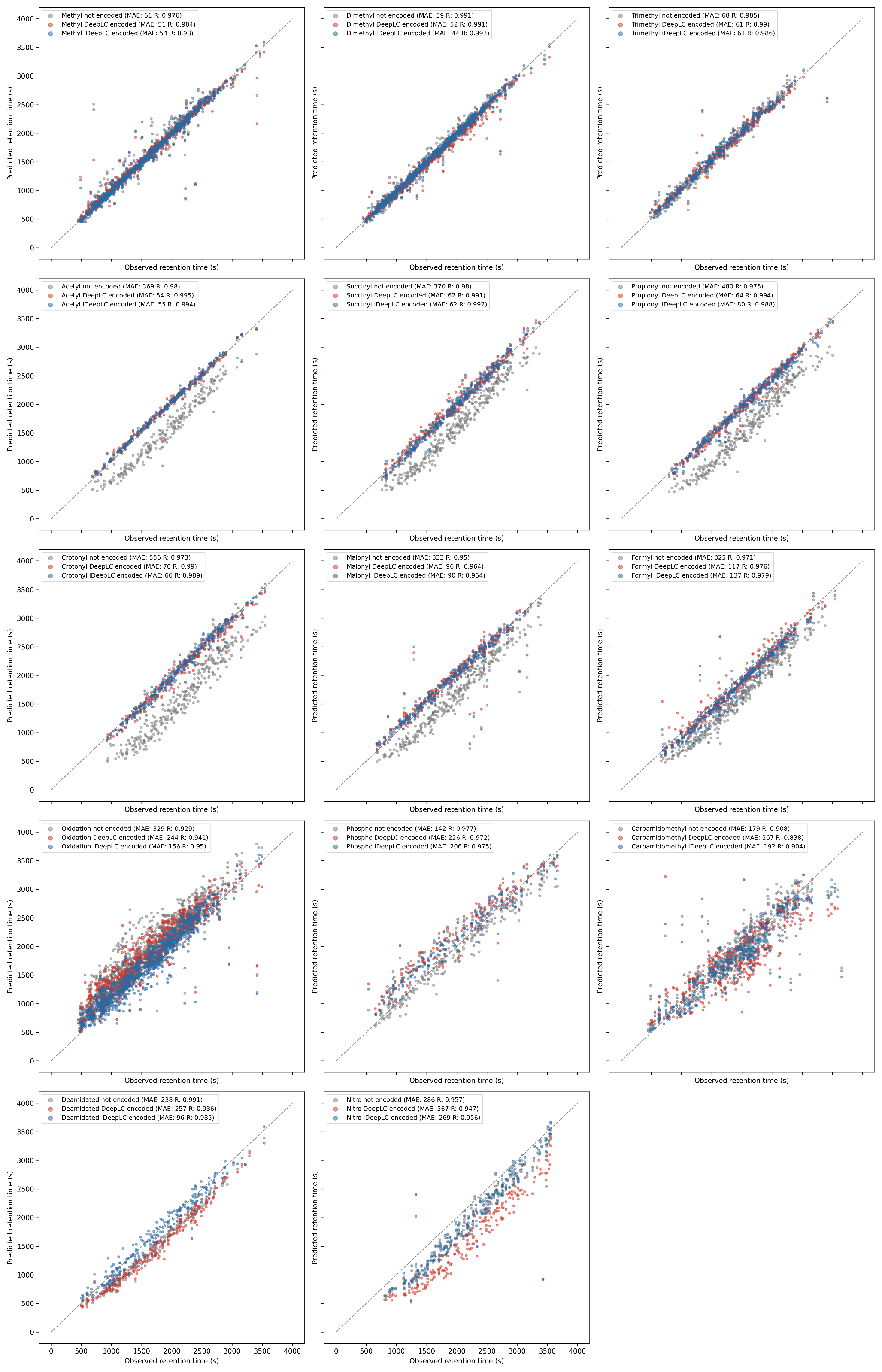

Supplementary Figure 1: Each subfigure shows the observed against predicted retention time for iDeepLC and DeepLC models not trained on one specific modification. The blue and red dots show the retention time on an iDeepLC model and a DeepLC model respectively trained with encoded modifications, and the grey dots show the retention time on a model trained with not encoded modifications. The variable MAE shows the error between predicted and actual retention time. And the variable R is the pearson correlation value between predicted and actual retention time.
